# Rapid eco-phenotypic feedbacks and the temperature response of biomass dynamics

**DOI:** 10.1101/2022.06.17.496633

**Authors:** Jean P. Gibert, Daniel J. Wieczynski, Ze-Yi Han, Andrea Yammine

## Abstract

As biomass dynamics capture information on population dynamics and ecosystem-level processes (e.g., changes in production over time), understanding how rising temperatures associated with global climate change influence biomass dynamics is a pressing issue in ecology. The total biomass of a species depends on its density and its average mass. Disentangling how biomass dynamics may respond to increasingly warm and variable temperatures may thus ultimately depend on understanding how temperature influences both density and mass dynamics. Here, we address this issue by keeping track of experimental microbial populations growing to carrying capacity for 15 days at two different temperatures in the presence and absence of temperature variability. We show that temperature influences biomass through its effects on density and mass dynamics, which have opposite effects on biomass and can offset each other. We also show that temperature variability influences biomass, but that effect is independent of any effects on density or mass dynamics. Last, we show that reciprocal effects of density and mass shift significantly across temperature regimes, suggesting that rapid and environment-dependent eco-phenotypic dynamics underlie biomass responses. Overall, our results connect temperature effects on population and phenotypic dynamics to explain how biomass responds to temperature regimes, thus shedding light on processes at play in cosmopolitan and massively abundant microbes as the world experiences increasingly hot and variable temperatures.

## INTRODUCTION

Understanding the biotic and abiotic factors that influence ecosystem function is a central goal of ecology (Giller and O’Donovan 2002, Srivastava and Vellend 2005, Begon et al. 2006). While censusing species presence/absence and abundances (or densities) provides a window into the overall structure of a community (e.g., composition, richness, evenness, diversity), species abundances alone do not contain information on the ecosystem-level functions performed by that community. However, tracking biomass (or biomass density) over time –i.e., the total mass of all individuals of a species or community (per unit area, if biomass density)– provides information on production within trophic levels, and comparing biomass across trophic levels can yield information on energy transfers within a food web (McKie and Malmqvist 2009, Trebilco et al. 2013, D’Alelio et al. 2016, Barneche et al. 2021). Because of that, biomass is a central concept that both describes the state of an ecosystem and provides information on ecosystem-level processes that influence overall function like production or energy transfers (Hatton et al. 2015).

As the planet warms, the structural and dynamical responses of ecosystems are reflected in changes in biomass (Kortsch et al. 2015, Ullah et al. 2018, Bartley et al. 2019, Gibert 2019, Barbour and Gibert 2021). For example, the biomasses of multiple taxa have been shown to decline with temperature across systems (O’Connor et al. 2009, Carr et al. 2018, Larjavaara et al. 2021). However, biomass declines are not universal (Lin et al. 2010) and the mechanisms through which warming influences species biomass are not well understood. Intuitively, because biomass is the sum of the mass of all individuals in a species, it is possible to decompose biomass into two main components: species’ average masses and species’ abundances (densities). Indeed, biomass is often estimated in the field as the product of the average mass of the individuals of a population and their abundance (or density). Consequently, any effects of temperature on biomass should, at their core, result from temperature effects on the abundance/density of a species or its average body size/mass.

Body size is an important functional trait that determines metabolic rates (Gillooly et al. 2001, Brown et al. 2004), demographic parameters (Savage et al. 2004, DeLong and Hanson 2009, Wieczynski et al. 2021), species interactions (DeLong 2014, DeLong et al. 2014, DeLong et al. 2015), and even community and ecosystem-level structure and processes (Allen et al. 2005, Gibert and DeLong 2014, Schramski et al. 2015, Wieczynski et al. 2021). Increasing temperature generally reduces individual body sizes, an effect called the ‘temperature-size rule’ (TSR) that is pervasive across systems and taxa (Atkinson 1994, Atkinson 1995, Atkinson et al. 2003, Forster et al. 2012). For these reasons, body size and the temperature-size rule have clear consequences for changes in biomass across all levels of ecological organization in a warming world (Brose et al. 2012).

How temperature influences the other component of biomass –density– is less clear. The Metabolic Theory of Ecology predicts that warming should decrease species’ carrying capacities–the maximum density attainable in a given environment– but proof of that decline remained elusive until recently. Data-tested theoretical work has now shown that carrying capacity indeed declines with temperature, but this effect can only be understood by integrating associated effects on body size via the TSR (Bernhardt et al. 2018). Moreover, while carrying capacities may indeed decline with temperature, it is unlikely that all species within a community will be at carrying capacity at any given moment—rather transient, non-equilibrium dynamics are expected (Hastings et al. 2018). Thus, addressing whether and how non-equilibrium densities are impacted by temperature is important for understanding how temperature influences biomass.

Last, body size can influence population growth, and hence densities, through relationships with demographic parameters like carrying capacity (K) and the intrinsic growth rate (r) (Damuth 1981, Savage et al. 2004, DeLong et al. 2015). On the flip side, population dynamics could, in theory, also influence body size, through associated effects on resource levels, but these effects are less well understood. Recent work has shown that, as populations grow to carrying capacity, rapid changes in body size can have a stronger effect on changes in density than the other way around, suggesting important –albeit asymmetric– feedbacks between population density and body size (Gibert et al. 2022). But how these reciprocal effects change with temperature, or how they may influence biomass responses to warming, is not known.

Here, we tackle these unknowns by addressing the following questions: 1) How is biomass affected by temperature and temperature variability as a species grows to carrying capacity? 2) To what extent are the effects of temperature on biomass dependent on how density and body size dynamics respond to temperature? 3) Does density or body size have a stronger effect on biomass responses to temperature? And, 4) do the reciprocal impacts of density and body size vary across temperature regimes? To address these questions, we recorded time series of population dynamics in a microbial species and tracked changes in total biomass, density, and body size in four different temperature regimes: constant 22ºC, constant 25ºC, and both temperatures with fluctuations. We derive a simple mathematical expression to partition the contribution of changes in density and body size to changes in biomass and assess how temperature responses in either one influence biomass shifts. Last, we use time series analyses to assess whether and how reciprocal effects of density and body size on biomass vary across temperature regimes.

## METHODS

### Microcosm growth assays

We grew populations of the protist *Tetrahymena pyriformis* for 15 days at various temperature treatments. To do so, we set up 24 experimental microcosms in 250 mL autoclaved borosilicate jars containing 200 mL of Carolina protist pellet media (1L of autoclaved DI water per pellet) previously inoculated with pond bacteria from Duke Forest (Gate 9/Wilbur pond, Lat=36.02°, Long=-78.99°, Durham, NC) and a wheat seed as a carbon source for the bacteria (Altermatt et al. 2015). All microcosms were started at 10 ind/mL protist densities and incubated in humidity-controlled (65% humidity) growth chambers (Percival AL-22L2, Percival Scientific, Perry, Iowa) on a 12hr night/day cycle. The entire replicated timeseries is therefore composed of 360 data points.

The 24 microcosms were subdivided into 4 experimental treatments: constant 22ºC, constant 25ºC, variable 22ºC or variable 25ºC. Temperature variability was programmed into our growth chambers to keep an average temperature of either 22ºC or 25ºC, but to fluctuate between two temperatures that were ±1.5ºC of the average every 12 hours, therefore imposing variability with a thermal amplitude of 3 ºC. A microcosm in the variable 22ºC treatment thus spent half of the day at 19.5ºC and half of the day at 23.5ºC while one in the variable 25ºC spent half of the day at 23.5ºC and half of the day at 26.5ºC. At each temperature change, temperature ramped up/down for roughly 15 minutes. Neither water nor nutrients were replaced throughout the course of this experiment. From now on we call these temperature treatments constant’ (C) and ‘variable’ (V).

### Density, mass, and biomass estimates

Densities (ind/mL) and trait dynamics were tracked daily for 15 days through fluid imaging of 1 mL subsamples of each microcosm (Fig 1a, FlowCam, Fluid Imaging Technologies, Scarborough, ME, USA). The FlowCam captures images of particles ranging from 5-10 μm to 2mm in length. The procedure produced ∼ 450k cell images, thus providing us a unique window into how biomass, density, and body size, changed together over the course of this experiment. Density was quantified as cell counts per volume sampled. Cell mass was quantified as the product of cell volume (as the volume of a spheroid, in μm^3^) and water density (1 g/cm^3^, or 10-12 g/μm^3^). Sample biomass was measured as the sum of the masses of all individuals per sample (in grams, g). However, the FlowCam can only census a fraction of each water sample. This determines the efficiency of the machine (in our case ∼ 0.33). Because of that, total biomass needs to be corrected by the efficiency, as the observed number of individuals is a fraction of the total that actually occur in our water samples. To do so, we linearly transform sample biomass according to the observed relationship between the number of cell images and the actual densities as detailed in Appendix 1. True biomass is therefore the observed biomass divided by the sampling efficiency.

**Fig 1:**
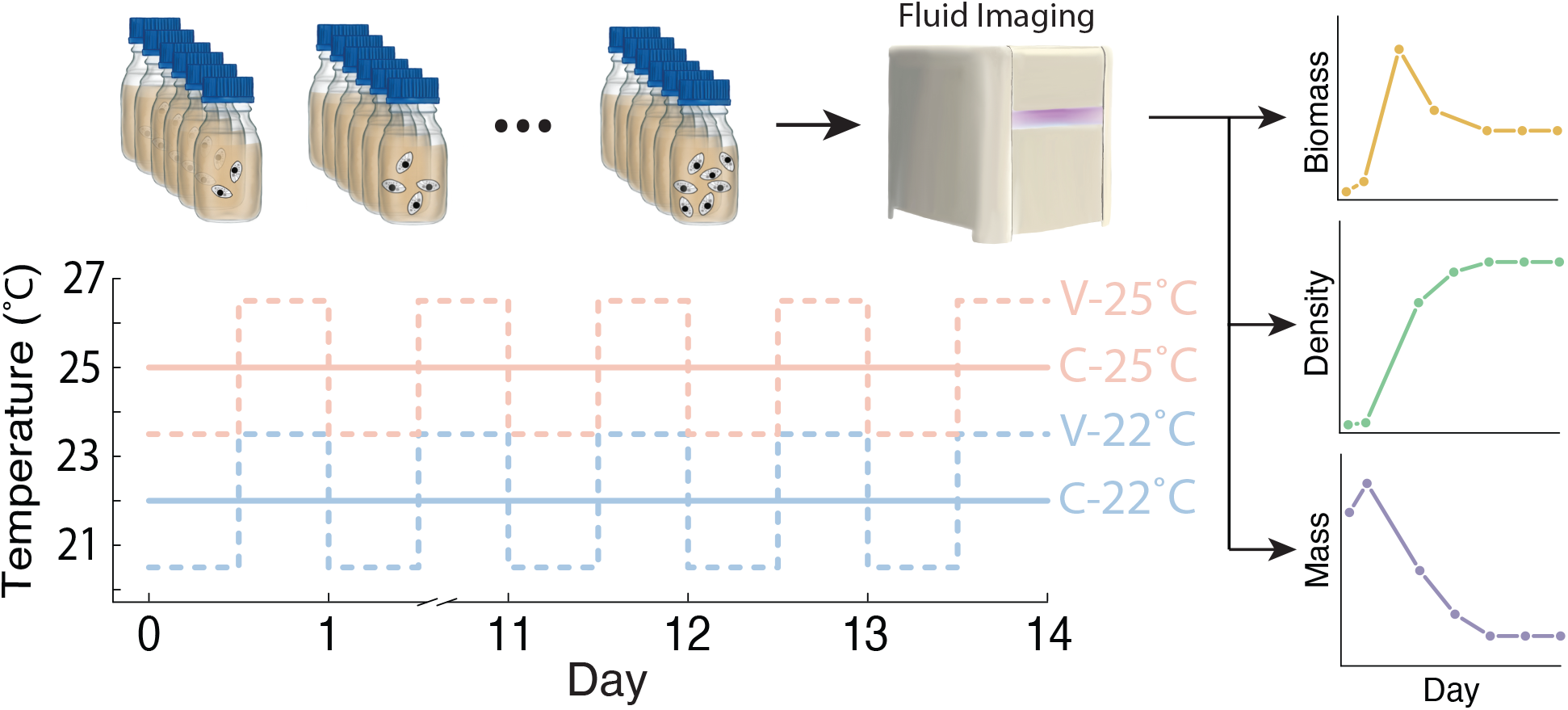
Microcosms where initialized at Day 0 and kept in four possible temperature treatments (Constant-22ºC, Variable-22ºC, Constant-25ºC or Variable-25ºC) for 15 days. Each day, a 1mL sample of media was taken for fluid imaging (FlowCam) to estimate total biomass, density, and average mass as the species grew to carrying capacity.

### Statistical analyses

To assess how temperature regimes influenced biomass, density and mass dynamics, we used Generalized Additive Mixed Models (GAMMs) with biomass, density, or mass as the response variables, day is a smooth term, both temperature and the presence and absence of variation as discrete predictors, and jar replicate as a random intercept. Additionally, because time series are necessarily sampled in a repeated fashion within each replicate, temporal autocorrelation may exist. To account for this temporal autocorrelation, we included an Autoregressive Moving Average (ARMA) correlation structure of order one in our GAMMs using the R package mgcv v.1.8 (Wood 2011, Wood et al. 2016).

While GAMM yields a good understanding of how time and treatments influence dynamics, a finer understanding is possible by assessing what specific aspects of the dynamics may have been influenced by the treatments. First, we assessed whether the imposed treatments in any way influenced the peak observed biomass by running a multiple linear regression (‘lm’ function in base R (R Core Team 2013)) with peak biomass (i.e., from days 3 to 5) as the response variable and both additive and interactive effects of temperature and the presence/absence of fluctuations as predictors. To quantify which differences between treatments were significant, we also ran a separate ANOVA with a post-hoc Tukey test (‘aov’ and ‘TukeyHSD’ functions in base R (R Core Team 2013)) with peak biomass (i.e., from days 3 to 5) as the response variable and all four temperature treatments as separate predictors. We used the same statistical methods to assess whether demographic parameters controlling density –i.e., intrinsic growth rates, r, and carrying capacities, K– changed with treatment. Intrinsic growth rates *r* where calculated as the natural log of the ratio of the density at day 1 and the density at day 0 (Wieczynski et al. 2021, Gibert et al. 2022), and K was estimated as the densities measured over the last 2 days of the dynamics in each jar.

### Decomposing change in biomass into change in density and mass

To decompose the contribution of changes in density and mass to the observed changes in biomass, we assume that the biomass, B, could be written as a function of density, N, and average mass, M, as

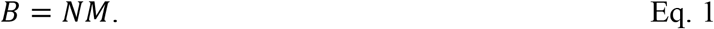

The rate of change of B over time, 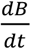, can be found by taking time derivatives in both sides of Eq. 1., which yields:

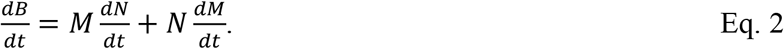

We then noticed that Eq 1 could be used to solve for either N or M, as 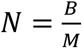 and 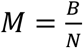, and replaced both into Eq. 2 to get:

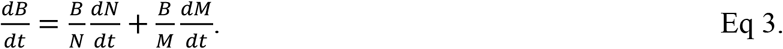

Eq. 3 coul be further simplified by factoring B, dividing both sides of the expression by B, then using the relation 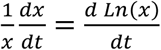 to get:

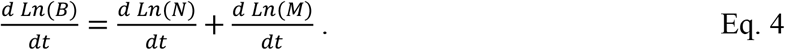

Eq. 4 links the rate of change in Ln(B), to that of Ln(N) and Ln(M). This equation can thus be used to decompose the contributions of N (i.e., 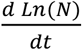) and M (i.e., 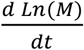) to the rate of change in B over time and across temperature treatments. We used our experimental time series to calculate these contributions of N and M to changes in B for each individual jar on each day of the experiment.

### Time series analysis

Previous studies have shown that Convergent Cross Mapping (CCM) can be used to infer causation between variables with available time series across ecological systems and environmental conditions (Sugihara et al. 2012, Clark et al. 2015, Karakoç et al. 2020, Kondoh et al. 2020, Rogers et al. 2020). A recent study used CCM to show that rapid plastic change in body size influences population dynamics more than the other way around, which was then confirmed through a manipulative experiment (Gibert et al. 2022). Following this literature, we therefore used CCM to assess whether change in body size more strongly influenced changes in density, or vice versa, across the temperature treatments.

CCM quantifies whether one time series (A) causally influences another (B) through the estimation of how much information of A is contained in B (Takens 1981, Sugihara et al. 2012). Conceptually, if a variable A causally influences variable B, but B does not influence A, then B should contain information about A, but not the other way around. CCM assesses how much information of the one variable is contained in the other by quantifying whether variable A can be predicted from the time series of B (and vice-versa) for subsets of the time series of increasing length (the length of these re-sampled time series is called the library size). If A more strongly influences changes in B than the other way around, then B responds to A more strongly than A responds to B. If the effect of A on B is causal, then the ability to predict A from B increases with library size, while the error associated with the prediction decreases. If this ‘predictability’ (or cross-mapping skill, ρ) is constant across library sizes, there is correlation, but not causation (Sugihara et al. 2012). More details can be found in the now extensive literature on this algorithm (Brookshire and Weaver 2015, Ye et al. 2015a, Ye et al. 2015b, Kaminski et al. 2016, Hannisdal et al. 2017, Luo et al. 2017, Mønster et al. 2017, Tsonis et al. 2018, Vannitsem and Ekelmans 2018, Liu et al. 2019, Barraquand et al. 2020). We used modified version of the CCM algorithm (R package multispatialCCM v1.0 (Clark et al. 2015)) to analyze the time series for each of the four temperature treatments because it allows for replicated times series.

## RESULTS

### General dynamics

Biomass increased steeply in the early days of the dynamics, then declined over time (Fig 2) across temperatures. Density showed a typical logistic growth pattern of fast growth in the early days followed by a plateau at around 6,000 ind/mL (Fig 2b). Average mass increased from Day 0 to Day 1, then decreased roughly monotonically over time (Fig 2c).

**Fig 2:**
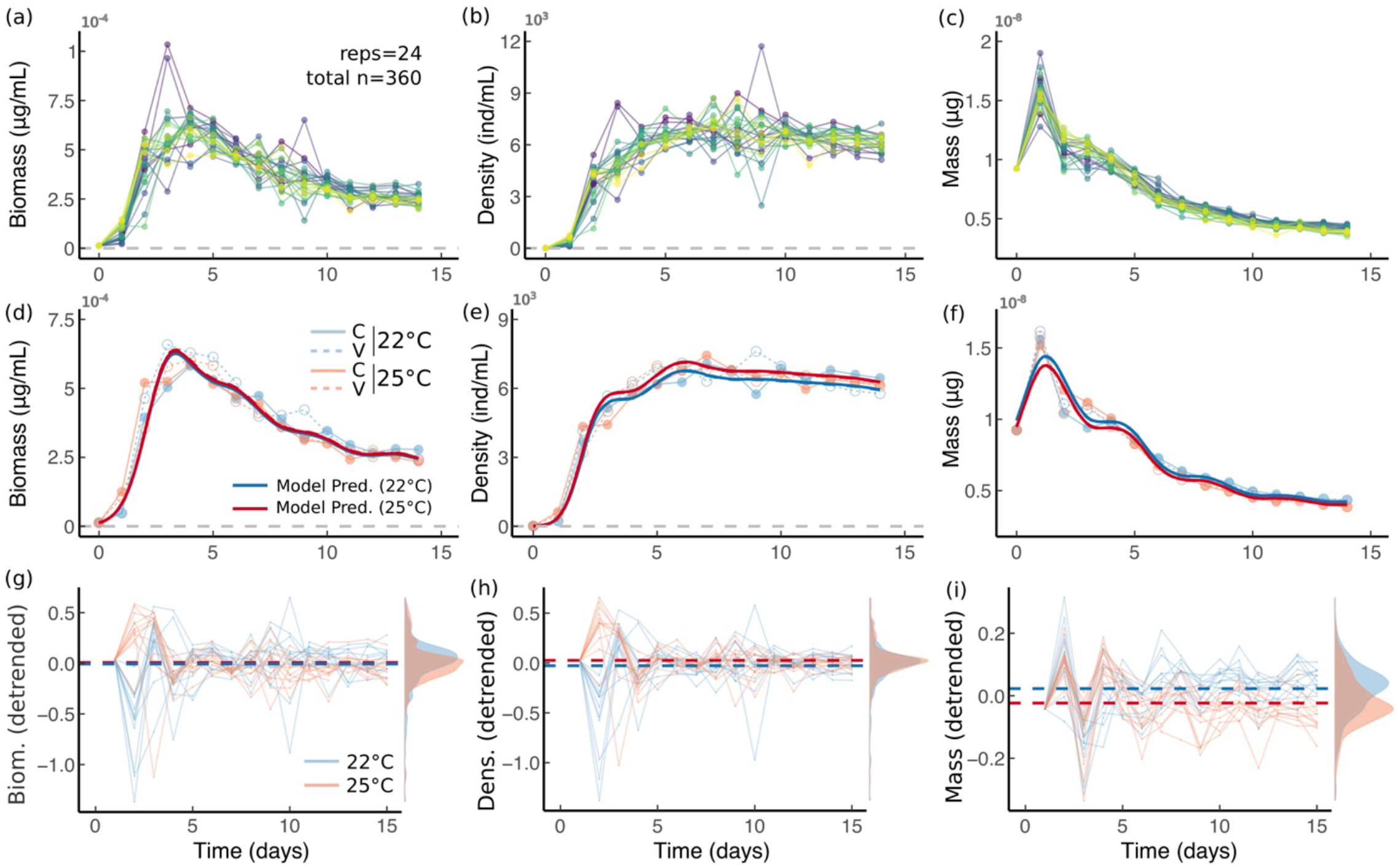
(a) Biomass over time for all 24 experimental jars. (b)-(c) as in (a) but for density and average mass respectively. (d) Biomass change over time (dots represent average biomass across all 6 replicates within each experiment, blue represents 22ºC treatments, red represent 25ºC treatments, while solid lines represent constant temperature treatments, C, dashed lines represent variable temperature treatments, V). Solid bold lines represent GAMM model predictions. (e-f) as in (d) but for density and mass. (g) Detrended biomass dynamics across all temperature treatments (but only color coded for mean temperature as temperature variability had no effect in d-f) with color coding as in (d). Bold dashed lines represent mean biomass for both temperature treatments. The distribution that biomass values take over time are shown on the right. (h-i) as in (g) but for density and mass.

### Effects of temperature and variability on biomass, density, and average mass

Biomass did not respond to either temperature (estimate=0.02±0.02, p=0.48, Fig 2d) or temperature variability (estimate=-0.009±0.02, p=0.73, Fig 2d). Temperature had a positive additive effect on density at 25ºC relative to 22ºC (estimate=0.05±0.02, p=0.018) while temperature variability had no effect (estimate=0.003±0.02, p=0.89, Fig 2e). Temperature also had a negative effect on mass (estimate=-0.006±0.003, p=0.002), but there was no effect of variability (estimate=-0.03±0.01, p=0.06, Fig 2f). These results suggest that the effects of temperature on density and mass likely cancel each other out, thus leading to an apparent lack of biomass temperature response.

Once the time series were detrended (by subtracting a GAMM model only containing time as a smooth term), additional effects of the treatments could be observed (Fig 2 g-i). In particular, biomass and density showed similar strong effects of temperature (but not fluctuations) in the first few days of the dynamics (Fig 2g & 2h). Mass temperature responses, however, were most prevalent in the later dynamics, when the temperature size rule appears to set in (Fig 2i).

Despite showing no effects of temperature or variability on overall biomass dynamics (Fig 1d-f), peak biomass in the variable environment was higher than in the non-variable environment across temperatures, and this difference was only slightly smaller in the high temperature treatment, thus showing an effect of temperature variability but not temperature alone on peak biomass (temp. effect = 4×10^-7^±3×10^-6^, p=0.906, var. effect = 9.492×10^- 6^±3.410×10^-6^, p=0.007, interaction= -5×10^-6^±5×10^-6^, p=0.314; ANOVA p = 0.02, Fig 3a).

**Fig 3:**
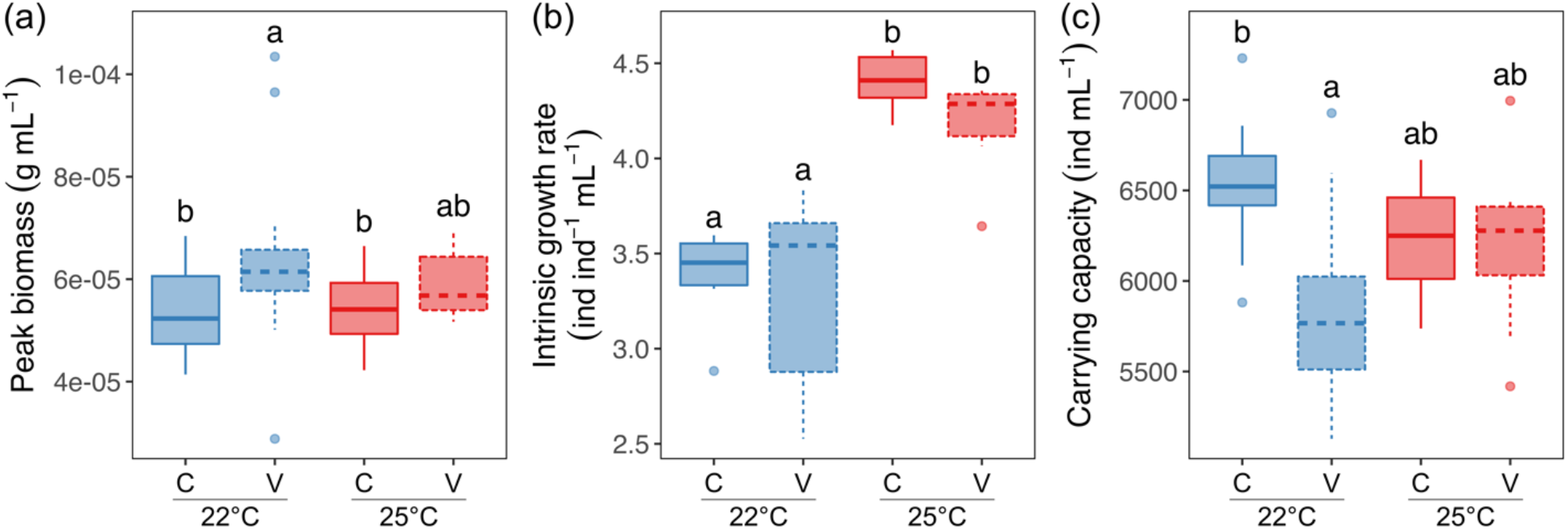
(a) Peak biomass, measured at days 3-5 with significant differences indicated as letters above the boxes. Variable temperatures lead to higher peak biomass, and that peak is higher at 22ºC than at 25ºC. (b) Intrinsic growth rate increases at 25ºC. (c) Carrying capacity is higher at constant 22ºC than in variable temperature but that difference disappears at 25ºC. Color coding as in Fig 1.

Temperature and temperature variability also influenced simple descriptors of what are otherwise complex density dynamics (Fig 3b, c). Indeed, temperature increased intrinsic growth rate despite fluctuations having no effect (temp. effect = 1.03±0.02, p<10^-4^, var. effect = -0.08±0.02, p=0.7, interaction= -0.17±0.28, p=0.6; ANOVA p < 10^-4^, Fig 3b; calculated using the first two days). Carrying capacity, on the other hand, decreased with variability but only at the low temperature and showed no significant differences between temperatures (temp. effect = -304±166.1, p=0.074, var. effect = -696±166, p<10^-3^, interaction= 667.2±234.9, p=0.007; ANOVA p = 0.002, Fig 3c).

### Decomposing the effects of density and mass on biomass across treatments

Density and mass dynamics contributed distinctly to biomass dynamics, especially in the first three days (Fig 4). For day ≤ 2, rapid density increases strongly and positively influenced biomass, while mass only positively influenced biomass dynamics on day 1, then made mostly negative contributions (GAMM smooth term = p<10^-16^, Fig 4a; ANOVA p<10^-16^, Fig 4g), likely due to the monotonous decline in mass from day 1 on (Fig 2c, f).

**Fig 4:**
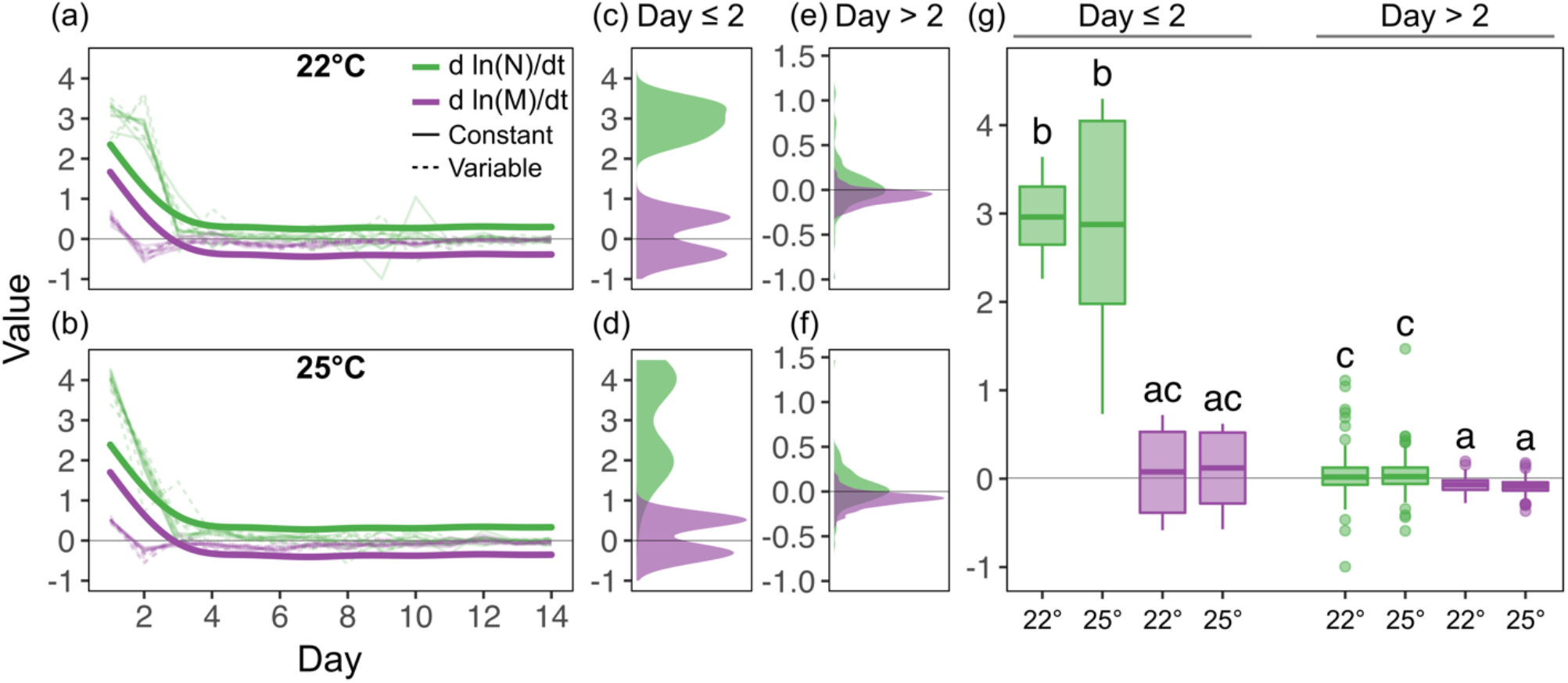
Contributions of density (d ln(N)/dt) and mass (d ln(M)/dt) to biomass dynamics at (a,c,e) 22ºC and (b,d,f) 25ºC. (a) and (b) show the contributions of density and mass over time at 22ºC and 25ºC, respectively. Thin lines represent replicate populations (jars) and thick lines are GAMM fits to these data. (c,d) and (e,f) show distributions of density and mass contributions over time for Days 0–2 and Days > 2, respectively. (g) shows differences in the contributions of density and mass across both temperatures for Day ≤ 2 and Day >2, evaluated using an ANOVA with Tukey’s HSD post-hoc test (p < 10^-5^).

Despite temperature and temperature variability influencing both density and mass dynamics, their effects on the contributions of either one to biomass dynamics –i.e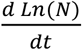 and 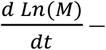 was surprisingly low, especially in the long-term. Initially (day ≤ 2), density had a large, positive affect on biomass that remained high until day 2 at 22ºC (Fig 4a) but declined sharply after day 1 at 25ºC (Fig 4b, thin lines). Beyond day 2, the contributions of either density or mass to biomass dynamics were small but different in sign (positive for density, negative for mass, Fig 4e–g). These results suggest that, while temperature treatment effects on biomass are most notable in the early dynamics, small, opposing effects of density and mass dynamics on biomass dynamics persist in the long term but are mostly unaffected by temperature and temperature fluctuations. Moreover, small temperature effects in the contributions of mass and density in the early dynamics are enough to produce larger effects later on.

### The temperature response of the coupling between density and mass

We observed that changes in mass more strongly influenced change in density than the other way around (consistent with a recent study (Gibert et al. 2022)) across all temperature treatments (Fig 5 and Fig S2 Appendix2). However, the strength of these reciprocal effects varied among treatments in specific ways. Temperature variability weakened the effect of mass on density across temperatures, and this effect was slightly stronger at 25 ºC compared to 22ºC (Fig 5, temp. effect = 0.013±0.001, p=0.15, var. effect = -0.05±0.009, p<10^-6^, interaction= -0.05±0.01, p<10^-4^). In contrast, the effect of density on mass weakened from 22ºC to 25ºC but got stronger with temperature fluctuations (Fig 5, temp. effect = -0.05±0.01, p<10^-6^, var. effect = 0.20±0.007, p<10^-16^, interaction= -0.02±0.01, p=0.29). These results suggest that rapid feedbacks between density and mass (or “eco-phenotypic feedbacks”) may themselves depend on environmental conditions—especially the effect of density on mass, which seems to respond more strongly to environmental variability than the effect of mass on density (Fig 5).

**Fig 5:**
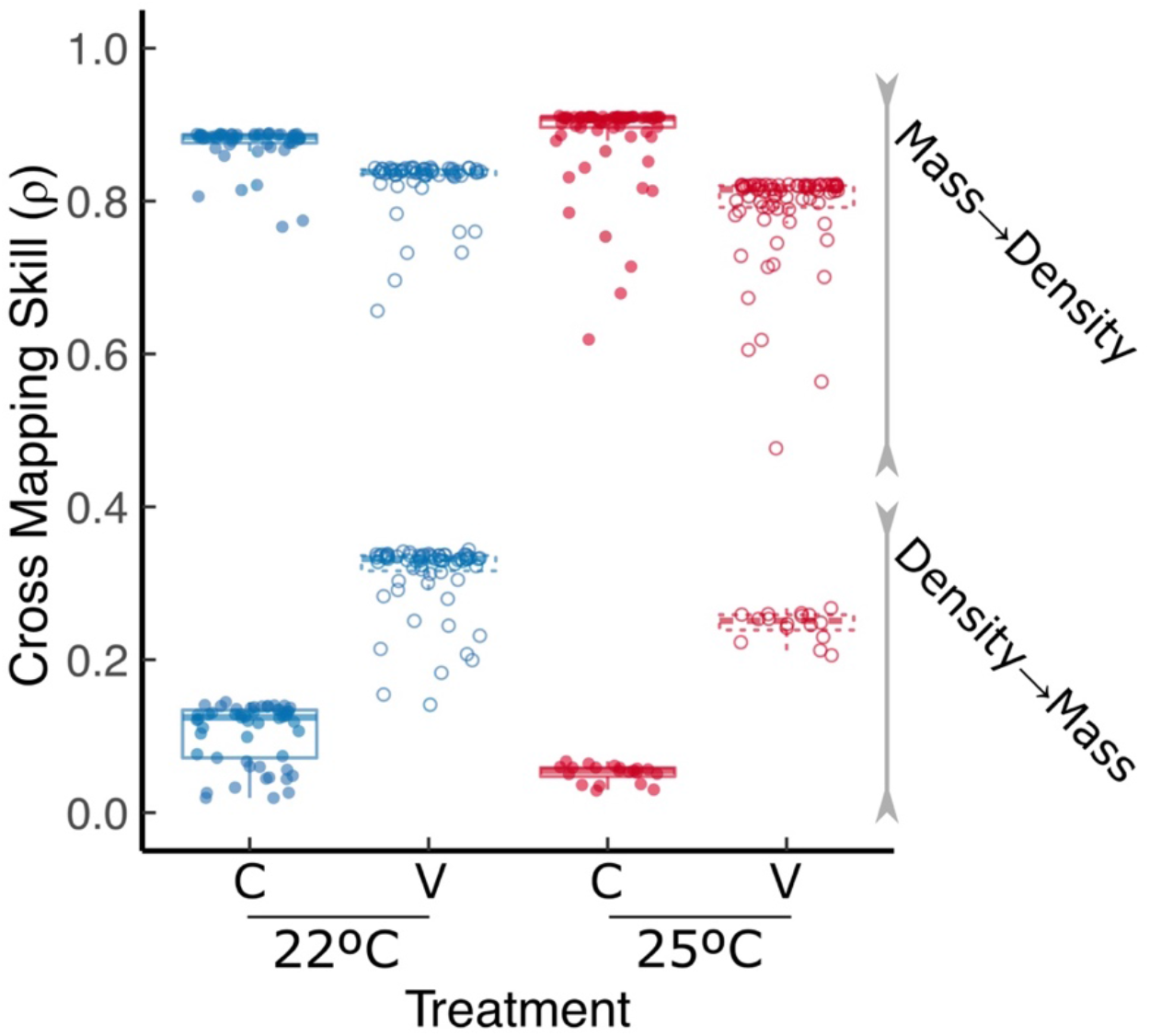
We show the cross-mapping skill (ρ) for all possible library sizes across temperature treatments (here represented as individual dots). Package multispatialCCM performs CCM on 800 total bootstrap replicates of the time series for each library size and yields an average value for the cross mapping skill. Effects of mass on density are indicated as ‘Mass→Density’ and effects of density on mass are indicated as ‘Density→Mass’. Mass more strongly influences density than the other way around, but the reciprocal effects of mass and density respond to both average temperature and temperature fluctuations. Colors as in Figs 1-3.

## DISCUSSION

Understanding how changes in environmental conditions influence biomass dynamics is paramount in ecology. Here, we argue that doing so requires understanding how temperature and temperature variability influence density and mass dynamics, then determining how those, in turn, influence biomass dynamics. Our results show that, while density and mass dynamics are independently susceptible to changes in temperature regimes (Fig 2 and 3), these effects may cancel each other out and not always translate to changes in biomass in response to temperature (Fig 2). We also show that different aspects of density-mass-biomass dynamics respond differentially to variation in environmental conditions (Fig 3), even when environmental effects on overall dynamics are less obvious (Fig 2). We show that density and mass have mostly opposite effects on biomass and their contributions are nuanced and likely stronger in earlier dynamics (Fig 4). Last, we show that temperature and temperature variability can alter the strength of feedbacks between mass and density (Fig 5), suggesting that rapid eco-phenotypic feedbacks may play and important but poorly understood role in biomass change in novel environments.

Previous research has shown that temperature often reduces body size, a phenomenon widely known as the Temperature-Size Rule (or TSR, e.g., (Atkinson 1994, Atkinson et al. 2003)). This phenomenon is widespread in myriad organisms, including mammals (Ozgul et al. 2009), birds (Weeks et al. 2020, Jirinec et al. 2021), invertebrates (Ghosh et al. 2013) and unicellular organisms (Atkinson et al. 2003, DeLong 2012, Tabi et al. 2020). The TSR has long been suggested to play an important role in the responses of populations (Ozgul et al. 2009), communities (Brose et al. 2012, Forster et al. 2012, Gibert and DeLong 2017) and ecosystems (Brose et al. 2012) to warming, as body size can directly impact reproductive and mortality rates and species-interaction parameters through its effect on metabolic rates (Gillooly et al. 2002, Brown et al. 2004, Savage et al. 2004). Our results show that the onset of the TSR occurs very early in population dynamics as species grow towards carrying capacity (Fig 1). Our results also suggest that, despite the numerous hypothesized effects of the TSR on ecological processes and dynamics, the TSR represents at most 5% of the observed variation in mass over time, with transient changes in mass being much larger in magnitude than the observed TSR (Fig 1).

However, recent work has shown that, without accounting for the TSR, predictions about how temperature influences long-term species densities (i.e., at carrying capacity) may be inaccurate (Bernhardt et al. 2018). Our results further imply that, without accounting for the TSR, inferring changes in biomass from changes in density alone may lead to equivocal estimates, as the effects of temperature on density and mass can cancel each other out (Fig 2). These results are important because they imply that environmental perturbations may – sometimes rapidly– change populations not just numerically (e.g., changes in densities), but also phenotypically. Although the ecological consequences of these rapid, plastic, phenotypic responses are still very poorly understood, our results emphasize the need to improve this understanding.

A recent study showed that rapid, plastic changes in body size more strongly influence changes in density than the other way around (Gibert et al. 2022), thus establishing the existence of important, but poorly understood, rapid feedbacks between body size and density. We observed the same pattern across in our study. Additionally, our results show that the strength of these feedbacks vary across temperature regimes (Fig 4) and that both mean temperature and temperature variability may be important. This result further emphasizes the need to study rapid phenotypic change –evolutionary or not– as a fundamental ecological response mediating how species cope with novel environmental conditions.

While our results provide novel insights about how rapid eco-phenotypic dynamics may mediate changes in biomass, density, and mass in response to warming and temperature variability, we caution against extrapolating these results beyond the range of temperatures studied here. Indeed, temperature effects are well known to have canonically unimodal effects on many demographic rates (Amarasekare and Savage 2012, Amarasekare and Coutinho 2013, Luhring and DeLong 2017, DeLong et al. 2018, Wieczynski et al. 2021). Thus, eco-phenotypic responses to a wider range of temperatures may be more complex than the results reported here. Moreover, the regimes of temperature fluctuations imposed here were less variable than the random fluctuations expected in an increasingly warmer world (Vasseur et al. 2014). Because of this, we also caution against interpreting our results to say that average temperatures cause stronger species-level responses than temperature variability and, in fact, some of our results even suggest that variability does have important effects (Fig 3a, Fig 5). Last, while CCM has long been used to infer effects of one time series on another (e.g. (Sugihara et al. 2012, Clark et al. 2015, Ye et al. 2015a, Ye et al. 2015b, Tsonis et al. 2018)), other unobserved variables like reductions in available nutrients, effects of regular sampling, or even physiological and metabolic changes as the ecological dynamics unfold, may affect and even weaken the CCM inference. A silver lining is that our results are consistent with those obtained by Gibert et al. (2022) which were validated with additional body size and density manipulations and showed that CCM correctly inferred reciprocal effects between size and density based only on their time series, as was done here (Fig 5).

Overall, our results shed light on how rapid eco-phenotypic dynamics in density and mass may influence how biomass responds to changes in temperature regimes. Our study emphasizes the need to consider rapid phenotypic change as an important –but poorly understood– mechanism through which organisms cope with changes in environmental conditions, with important implications for species responses to a rapidly changing and increasingly warm world.

## ACKNOWLEDGMENTS

AY, DJW, and JPG, were supported by a U.S. Department of Energy, Office of Science, Office of Biological and Environmental Research, Genomic Science Program Grant to JPG, under Award Number DE-SC0020362. The authors thank Lorelei Van Gorder without whom the collection of this dataset would not have been possible in its current form.

