## Appendix for "Rapid eco-phenotypic feedbacks and the temperature response of biomass dynamics"

INDEX

Appendix 1	………………………………………………………………………………………………………………….	2

Appendix 2	………………………………………………………………………………………………………………….	3

Appendix 1: Rescaling densities to estimate biomass

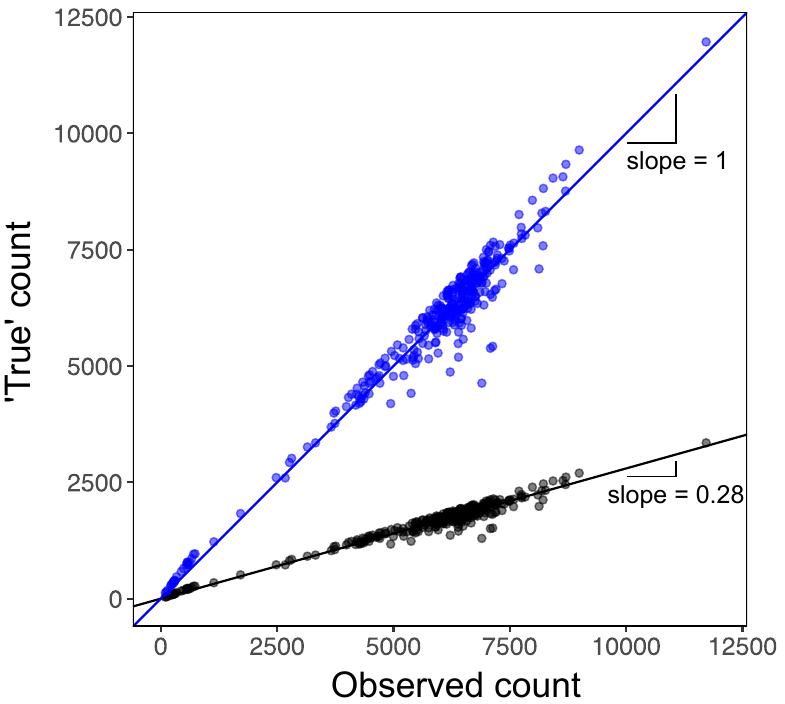

Fig S1: In black, the relationship between the observed number of individuals and the real number of individuals is shown before rescaling. In blue, the same relationship is shown after rescaling. True counts are estimated buy the FlowCam as the observed counts divided by the efficiency of the machine (in this case, the slope of the regression between observed and ‘true’, that is, 0.28). Each dot represents the observed and ‘true’ counts for each experimental jar for a given timepoint. Because total biomass is defined as the sum of the mass of all individuals, if the machine undercounts individuals, then we would be underestimating total biomass. To compensate for that, we estimate the actual biomass in our samples as the observed biomass divided by the efficiency (0.28). This rescaled biomass now accounts for the under-sampling of individuals and thus yields a more accurate estimate of biomass.

Appendix 2: Causation as inferred by Convergent Cross Mapping

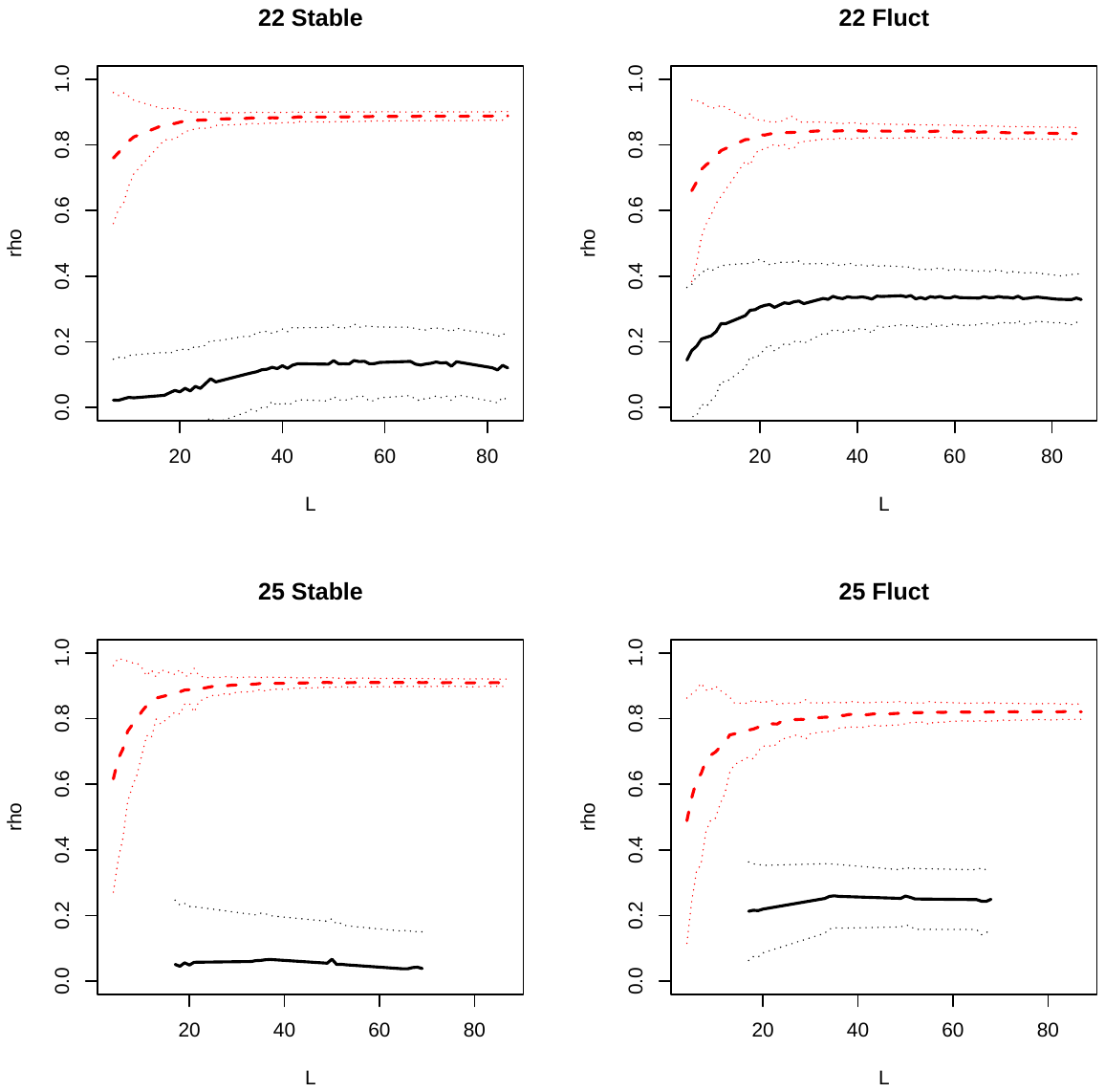
Fig S2: CCM results against library size for all four temperature treatments. Results indicate a strong causal effect of size on density (pink, dashed), and a weaker (sometimes non-causal or weakly causal) effect of density on size (black solid). Standard errors are shown as dotted lines. Results also emphasize that the reciprocal effect of size on density change with temperature regimes (e.g., effects of mass on density are slightly stronger in constant environments than in fluctuating ones while the effects of density on size seem more strongly causal [clear increase in rho with library size] at in the cold than in the warm).
